# Bioinformatics Copilot 2.0 for Transcriptomic Data Analysis

**DOI:** 10.1101/2024.08.15.607673

**Authors:** Yongheng Wang, Weidi Zhang, Ian Wong, Siyu Lin, Rohan Kumar, Aijun Wang

## Abstract

Large language models, when integrated into bioinformatics tools, offer valuable support for biologists in the analysis of single-cell transcriptomic data. Here, we introduce Bioinformatics Copilot 2.0, an advanced tool that builds upon its predecessor with four significant enhancements: 1) Internal server processing: users can now process large datasets on an internal server, thereby ensuring data privacy. 2) User-controlled analysis: users can take control of their data analysis when tasks exceed the copilot’s capabilities.3) Real-time information access: users can access up-to-date information from online resources, such as gene function queries and recent publications, thus enhancing data interpretation. 4) Data analysis and result documentation: the copilot now supports the generation of PDFs that include images and detailed analysis. These advancements represent a step towards building a bioinformatics autopilot. Demonstration videos showcasing these features are available at www.biochemml.com/tools.

## Introduction

In the past few years, there have been significant advancements in the field of single-cell spatial transcriptomics. These improvements enable the assessment of thousands of genes with subcellular resolution and provide unprecedented insights into cellular heterogeneity and the molecular distribution of tissues. For instance, NanoString has released 6,000-gene panels for its CosMx platform and plans to launch an 18,000-gene panel in 2025, allowing the evaluation of the transcriptome of human and mouse samples with a resolution of 200 nm. Similarly, 10X Genomics introduced 5,000-gene panels for its Xenium platform, also with a resolution of 200 nm for human and mouse tissues. Additionally, STOmics has introduced Stereo-seq OMNI, which makes it possible for full transcriptomic evaluation at a resolution of 500 nm for formalin-fixed paraffin-embedded (FFPE) tissue. The increase in gene profiling capabilities at high resolutions has significantly increased the complexity and volume of data produced. With the commercial availability of these advanced technologies, more labs can now access and utilize them, resulting in a rising demand for user-friendly analysis tools. Concurrently, the field of artificial intelligence has seen significant progress, with recent advances in prompt engineering^1^, agents^2^, retrieval-augmented generation (RAG)^3^, and fine-tuning^4^ and additional fields have enhanced the capabilities and broadened applications of large language models (LLMs). Techniques such as structuring LLM responses and retrieving information from additional databases can be effective in improving the performance of the Bioinformatics Copilot^5^. The parallel developments in spatial transcriptomics and AI offer a unique opportunity to leverage LLMs to address the analytical challenges posed by complex data.

## Result

Our updated application, Bioinformatic Copilot 2.0, introduces several new functionalities and an improved user interface compared to its predecessor. A key enhancement is the integration of a module that allows access to an internal server at the user’s organization (Figure 1), enabling them to log in and directly access server files. This is particularly beneficial for handling large datasets typical of single-cell spatial omics experiments, which can reach sizes of up to several terabytes or larger. Transferring data from an internal server to an external server can be time-consuming, and institutional policies often restrict sharing data externally, especially when patient samples and information are involved. Once logged into the server of a Linux system, users can simply right-click to download or rename files. This graphical interface is more intuitive than using command-line tools. The interface retains features from the previous version, such as fields for entering prompts and displaying work status or progress. Additionally, results will be displayed in the bottom right corner of the interface, with a scroll bar for easy navigation through reports (Fig. 1).

**Fig 1.**
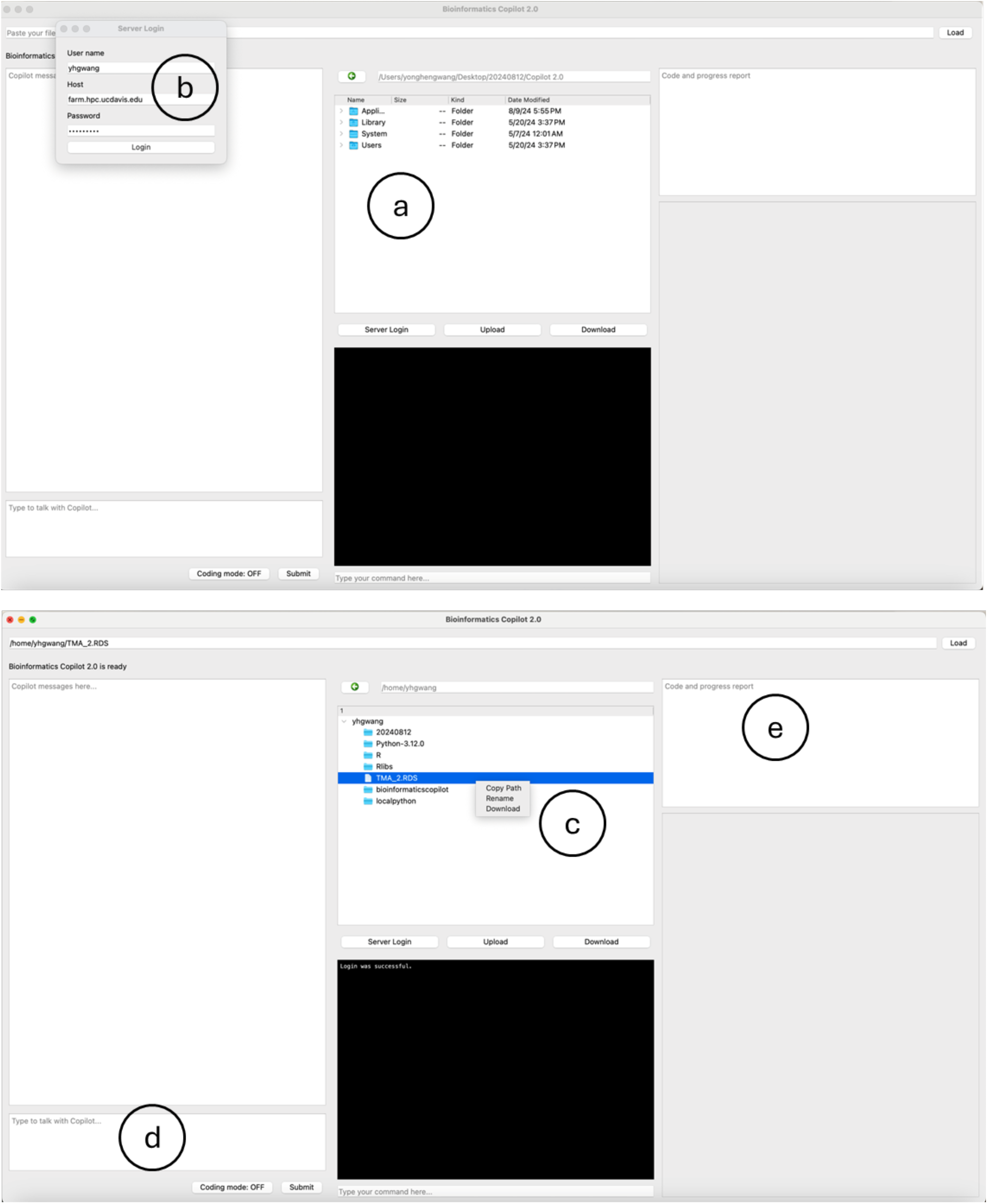
Software layout and functionalities. **a**, Panel for users to access files locally. **b**, Credential input area for connecting to a remote server. **c**, File management section for viewing, downloading, and renaming files on the server. **d**, Panel for users to enter commands and interact with the Copilot. **e**, Task monitoring area for tracking the status and progress of tasks.

The new version also broadens the spectrum of figure types that users can generate, including heatmaps, Kyoto Encyclopedia of Genes and Genomes (KEGG) pathway maps, and dimension plots (Fig. 2). Novice users can use the prompt and LLM feature on the left side of the software to request code for generating specific figures. Once the code is provided, users can copy and paste it to create the figures. For example, users can simply ask, “What is the code to make a heatmap in R?” and receive the code (Fig. 3b). Additionally, users can modify the software-generated code, such as adjusting parameters to change the size of dots in a dimensional plot. Experienced users can click the ‘Coding Mode’ button at the bottom left of the interface to code and debug independently. This feature can be useful for tasks that surpass the software’s built-in capabilities. If needed, users can ask the copilot for debugging advice using the tools on the left panel.

**Fig 2.**
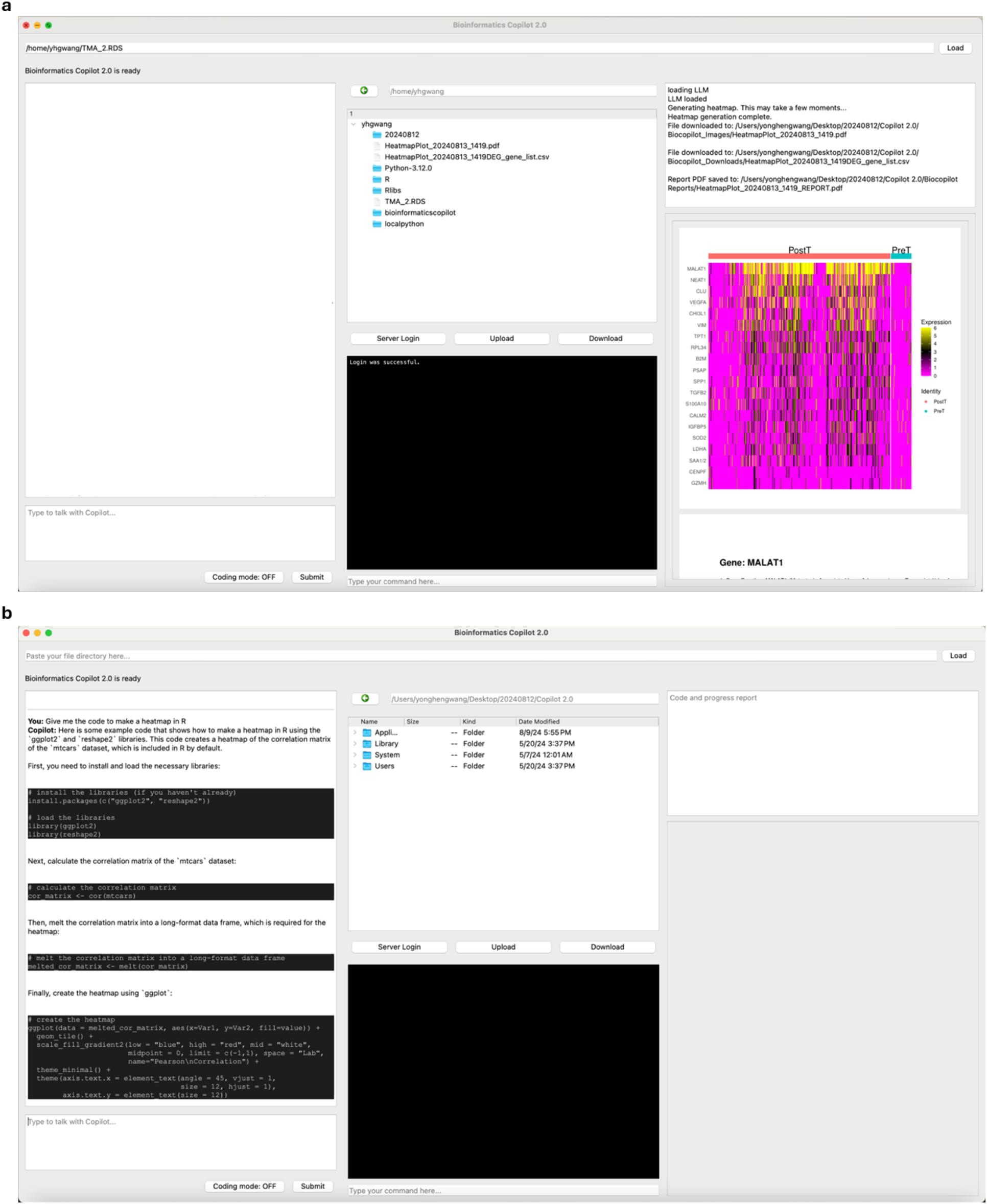
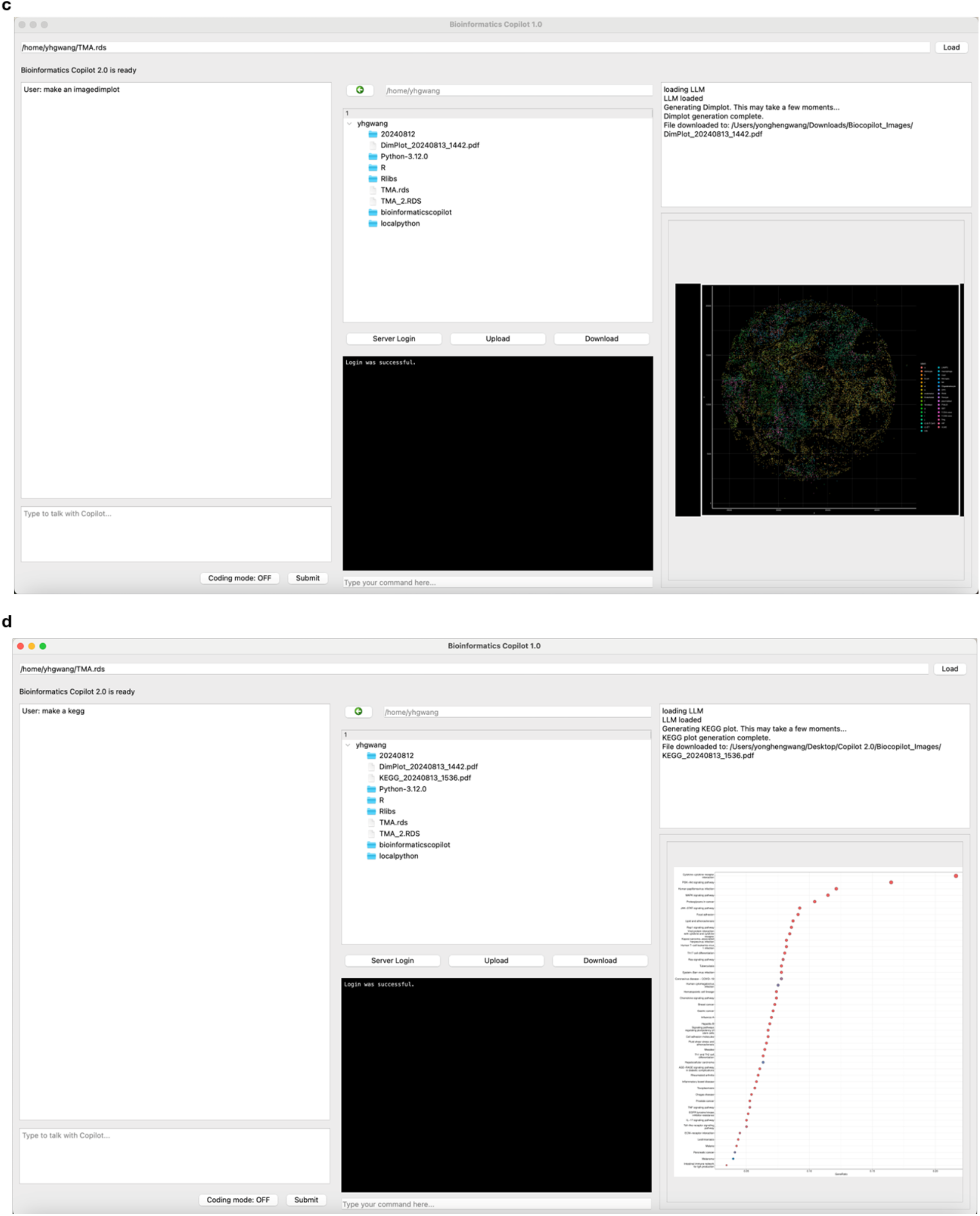
Result visualization and real-time interaction. **a**, Heat map displaying gene differential expression results derived from the transcriptomics data. **b**, Real-time query interface where users can obtain coding advice from the copilot **c**, Dimensionality reduction plot illustrating the clustering of the cells. **d**, Visualization of KEGG pathway enrichment results of the transcriptomics data, presented as a dot plot.

**Fig 3.**
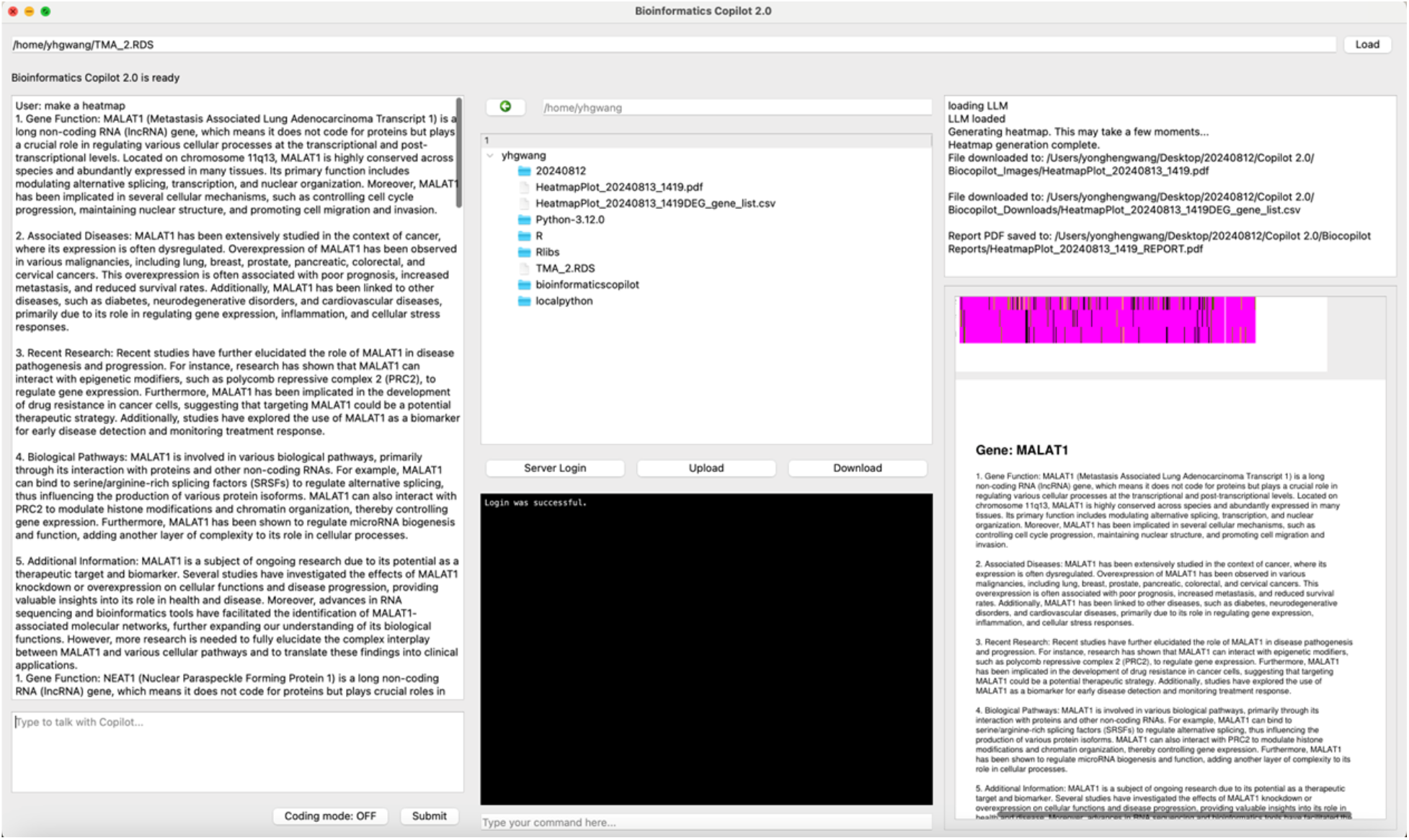
Automatic data interpretation and result summary. **a**, The software automatically analyzes the figures and retrieves relevant information from online sources. **b**, Figures and results are integrated into a comprehensive PDF report.

With permission, the software can automatically analyze data using the latest information, displaying the results on the left. For example, if users create a heatmap in which the top 5 differentially expressed genes are MALAT1, NEAT1, CLU, VEGFA, and CHI3L1, the software will perform a literature search and deeper analysis, presenting the findings on the left. Furthermore, the software can integrate the figures and analyses to create a comprehensive report, displaying it on the bottom right and automatically downloading it to the designated directory (Fig. 3). This feature represents a step toward creating a true bioinformatic autopilot.

## Discussion

Recent advancements in LLMs and generative AI have catalyzed the creation of new tools for deciphering transcriptomic data. For example, an attention-based machine learning model, Geneformer, has been introduced to predict chromatin dynamics and identify therapeutic targets^6^. Additionally, foundational models such as scFoundation^7^ and scGPT^8^, pretrained on extensive single-cell transcriptomic datasets, are being used to facilitate cell type annotation, perturbation prediction, and multi-omic integration. Various tools have been developed for omics data interpretation. OmicsAnalyst, for example, enables researchers to analyze complex multi-omics datasets without extensive programming or statistical expertise^9^. In contrast to OmicsAnalys, our software currently focuses specifically on single-cell transcriptomics and allows users to use natural language prompts and obtain updated online resources for debugging and data analysis. Moreover, many tools have been created for visualizing single-cell RNA-seq data, such as cellxgene, iSEE-loom, iSEE-SCE, loom-viewer, SCope, scSVA, Single Cell Explorer, and the UCSC Cell Browser. These tools mainly focus on visualization^10-11^, whereas our app is designed for data analysis using Seurat Objects as input.

The current version of our software faces several limitations, including restricted data format (Seurat objects only for the current version) and an insufficient depth of biological insights. To overcome these challenges, future iterations will incorporate additional modules to support a broader range of data types, such as H5AD and Fastq files. Leveraging RAG or pre-trained foundation models on biomedical literature will enhance the system’s ability to interpret data and generate meaningful biological insights. Key areas for improvement include bolstering the Copilot’s capabilities in suggesting research ideas, formulating hypotheses, and designing experiments^12^. Furthermore, the integration of more user-friendly features, particularly interactive visualization tools can make data analysis more intuitive for researchers.

## Notes

### Competing Interest Statement

The authors have declared no competing interest.

